# The peptide GOLVEN10 controls nodule and lateral root organogenesis and positioning along the longitudinal root axis

**DOI:** 10.1101/2022.05.06.490929

**Authors:** Sonali Roy, Ivone Torres-Jerez, Shulan Zhang, Wei Liu, Katharina Schiessl, Clarissa Boschiero, Hee-Kyung Lee, Patrick X. Zhao, Jeremy D. Murray, Giles E. D. Oldroyd, Wolf-Rüdiger Scheible, Michael Udvardi

## Abstract

GLV/RGF peptide encoding genes can be identified in genomes of all plants that can form roots or root-like structures suggesting they were essential for transition of plants to land.In *Medicago truncatula*, five of fifteen GOLVEN(GLV)/ROOT MERISTEM GROWTH FACTOR (RGF) peptide coding genes were induced during nodule organogenesis and to a varying extent under nitrogen deficiency and auxin treatment. Expression of *MtGLV9* and *MtGLV10* at nodule initiation sites was dependent on the transcription factor NODULE INCEPTION.Overexpression of all five nodule-induced *GLV* genes in *M. truncatula* hairy roots as well as application of the corresponding synthetic peptides resulted in a 25-50% reduction in nodule number indicating GOLVENs are negative regulators of nodule organogenesis.The peptide GOLVEN10 shifted the position of the first formed lateral root (rhizotaxis) as well as the first formed nodule along the longitudinal primary root axis, a phenomenon we term ‘nodulotaxis’, thereby reducing the absolute length of the zone of lateral organ formation on roots.Application of synthetic GOLVEN10 peptide caused an increase in cell number but not cell length in each root cortical cell layer causing an increase in root length and a consequent spatiotemporal delay in formation of the first lateral organ.

**Plain Language Summary:** Nodule positioning is an understudied trait, yet it determines the length of the root that can support nodule formation and consequently the total number of functional nodules formed. We identify for the first time, genetic factors called GOLVEN peptides that alter nodule and lateral root positioning on the primary root along with several other traits including nodule organ initiation and root architecture.

## INTRODUCTION

Root architectural responses to environmental cues such as availability of the macronutrient Nitrogen (N), are known to be complex and there is much to learn about the mechanisms that control root plasticity (Lynch, 2019). Changes in root system architecture involve priming, initiation and emergence of lateral organs that enable plants to respond dynamically to fluctuations in supply and demand of water and nutrients (Laskowski & ten Tusscher, 2017). In legumes, the ability to form nodules, a second type of root lateral organ, which in association with rhizobia help convert atmospheric-N into plant usable ammonia, adds yet another layer of complexity (Roy *et al*., 2020).

Plant hormones, including peptide hormones, are major determinants of root plasticity that bring about their effects by synergistic and antagonistic interaction between signaling networks (Matsuzaki, Yo *et al*., 2010; Meng *et al*., 2012; Whitford *et al*., 2012; Leyser, 2018; Zhu *et al*., 2020; Roy & Muller, 2022). Peptide hormones are short chains of amino acids that upon binding with their cognate cell surface receptors initiate a signal relay that ultimately controls physiological responses (Roy & Muller, 2022). The biological activity of chemically synthesized peptides predicted from genome sequences has revolutionized the field of chemical genomics. This has led to the discovery of multiple novel peptide hormones and helped uncover their roles in plant growth and root development (Okuda *et al*., 2009; Matsuzaki, Y. *et al*., 2010; de Bang *et al*., 2017). Two well-known families of peptide hormones, namely CLAVATA3/ENDOSPERM SURROUNDING REGION (MtCLE12, MtCLE13, MtCLE34, MtCLE35) and C-terminally ENCODED PEPTIDE (MtCEP1, MtCEP7), control lateral root and nodule development in *Medicago truncatula* (Mortier *et al*., 2010; Imin *et al*., 2013; Mens *et al*., 2021; Moreau *et al*., 2021). CEPs act as root-to-shoot ‘N-hunger’ signals controlling lateral root development and uptake of nitrogen in N-poor soils, while CLE peptides are perceived by the LEUCINE RICH REPEAT-RECEPTOR LIKE KINASE (LRR-RLK) receptor, SUPER NUMERIC NODULATION (SUNN) as part of a long-distance negative feedback loop called Autoregulation of Nodulation (AON) that restricts nodulation under N-replete conditions (Okamoto *et al*., 2013; Roy *et al*., 2020). Together these two pathways maintain an optimal N-balance and control nodule number in legumes (Laffont *et al*., 2020). Members of two additional families, PHYTOSULFOKINE (LjPSK8) and GOLVEN/ROOT GROWTH FACTOR/CLAVATA3 EMBRYO SURROUNDING REGION LIKE (MtGLV9/MtRGF3) have also been implicated as positive and negative regulators of nodule formation, respectively (Wang *et al*., 2015; Li *et al*., 2020).

The sulfated GOLVEN (GLV) peptides are known to control five root growth traits: cell number at the root meristem via activation of PLETHORA (PLT) transcription factors, which affects primary root growth (Matsuzaki, Y. *et al*., 2010); cell division during lateral root initiation thereby controlling lateral root density and patterning (Meng *et al*., 2012; Fernandez *et al*., 2015; Fernandez *et al*., 2020); auxin distribution via modulation of PIN efflux transporters thus regulating root gravitropism (Whitford *et al*., 2012); root cell number and circumferential cell growth rate under phosphate deficiency (Cederholm & Benfey, 2015); and nodule number (Li *et al*., 2020). At the molecular level, GLVs affect components of the auxin efflux transport pathway (PIN2) and auxin signaling (ARF7, ARF19) pathway and may reduce auxin concentrations at lateral root (LR) initiation sites (Whitford *et al*., 2012; Fernandez *et al*., 2020). Both nodule and lateral root formation are conditioned by changes in auxin flux and localized biosynthesis that lead to formation of local auxin maxima conducive for first cell divisions, organ initiation and outgrowth (Laskowski & ten Tusscher, 2017; Leyser, 2018). During nodule formation, this auxin buildup is associated with the transcription factor NODULE INCEPTION (NIN) (Schiessl *et al*., 2019).

Chemically synthesized plant hormones such as Indole-3-Acetic acid (3-IAA) and 6- Benzyl-Aminopurine (6-BAP) are instrumental in elucidating physiological roles of the classical hormones auxin and cytokinin, respectively (Michniewicz *et al*., 2019). Chemical genetics and synthetic peptides are equally valuable for understanding the role of peptide hormones in plant responses to their environment. Notably, discovery of LURE peptide, CEP peptide, and CLE peptide activity were all facilitated by their synthetic counterparts (Okuda *et al*., 2009; Goto *et al*., 2011; Corcilius *et al*., 2017; Laffont *et al*., 2020). The model legume *M. truncatula* has over 1800 potential genome encoded peptides, of which less than twenty have been functionally characterized (de Bang *et al*., 2017; Boschiero *et al*., 2020). Research on symbiotic nitrogen fixation in legumes over the past 20 years has focused on three main areas, including rhizobial infection (structure and number of infection threads), nodule organogenesis (morphology and number of nodules) and N-fixation rates (measured by acetylene reduction assays) (Roy *et al*., 2020). These processes have been investigated through forward and reverse genetics. An understudied aspect of root nodule symbiosis is control of nodule positioning. Penmetsa et al., observed that the position of nodule development relative to xylem and phloem poles were altered in Medicago *sickle* and *sunn* mutants (Penmetsa *et al*., 2003). Another hypernodulation mutant, *Ljplenty*, exhibited a wider zone along the primary root over which nodules were formed (Yoshida *et al*., 2010). In the current study, we investigated the biological role of nodule-induced GLVs and found that they decrease the length of the nodulation zone by altering nodule positioning along the primary root axis. We term this phenomenon ‘nodulotaxis’, by analogy to the term ‘rhizotaxis’ thatdescribes the arrangement or positioning of lateral roots along the primary root, which we also find to be under the influence of the GLV10 peptide.

## MATERIALS AND METHODS

### Plant material, nodulation assays and growth conditions

*Medicago truncatula* ecotype Jemalong A17 or R108 were used in this study. All *Tnt1* mutants are in the R108 background. *Tnt1* lines isolated include *glv10-1* (NF12742), and *glv10-2* (NF20983). Seeds were scarified with concentrated H_2_SO_4_, and surface sterilized with undiluted household bleach (Clorox) at 8% sodium hypochlorite and sown on 1% water agarose (Life Technologies Catalog: 16500100) plates.

Seeds were stratified at 4ºC for three days in dark prior to overnight germination at 24°C and then transferred onto agarose plates containing B&D nutrients (Broughton & Dilworth, 1971) plus 0.5 mM KNO_3_ with and without peptides. Eight to ten seedlings per line were placed on each plate between sterile filter paper sheets for all experiments except the GWAS screen in which we placed three seedlings per plate. For experiments in soil, overnight germinated seedlings were transferred to a 2:1 mixture of turface:vermiculite. Plants were watered B&D solution containing six millimolar nitrogen before inoculation with rhizobia at seven days post germination. Post Inoculation, plants were subsequently watering with B&D nutrient media supplemented with 0.5 Mm Potassium Nitrate. All experiments were conducted in a controlled environment chamber at 24ºC under 16 hours light, eight hours dark conditions.

*Arabidopsis thaliana* wild type Col-0 or mutant lines were sterilized using bleach and 70% ethanol. After stratification for 3 days at 4º C, seeds were placed onto ½ MS (Murashige & Skoog) media plates with or without peptide and allowed to grow for 14 days. All plants were scored on the same day under a 4x Leica S7 microscope.

### Cloning of promoters and CDS for hairy root

Gene coding regions were cloned either using Golden gate technology or the Gibson assembly method. Promoter regions of *MtGLV1* (1392 bps), *MtGLV2* (2883 bps), *MtGLV6* (2131 bps), *MtGLV9* (2192 bps), were cloned upstream of the β- Galactouronidase (GUS) gene in MU06 vector carrying a DsRed selection cassette using Gibson assembly method (Gibson *et al*., 2009). The *MtGLV10* endogenous promoter was synthesized (3000 bps) and cloned upstream of the GUS gene using Golden Gate cloning. All clones were verified by sanger sequencing before transformation into *M. truncatula* hairy roots mediated by *Agrobacterium rhizogenes* Arqua1 (Quandt *et al*., 1993). All cloned CDS using golden gate cloning (MtGLV1, MtGLV9, MtGLV10) and Gibson cloning (MtGLV2, MtGLV6) were cloned downstream of the Lotus *UBIQUITIN* promoter. Sequenced clones are available from Addgene under the deposit ID 79021.

### Hairy root transformation

A streptomycin resistant strain of *A. rhizogenes* Arqua1 was transformed with constructs of interest carrying a *AtUBI:dsred* selection marker and cultured on LB medium agar plates supplemented with the corresponding antibiotics for two days at 28°C, Agrobacteria were scraped off the plates using sterile spreaders and resuspended using 500-700 μL of sterile water. Root tips from overnight germinated seedlings were cut off to ensure that the meristem was completely removed and the cut end dipped in aforementioned bacterial suspension. Seedlings were transferred to and grown on modified Fahraeus medium plates for two weeks at 24ºC 16 hours day and 8 hours night conditions. Transgenic calli expressing dsRed were selected using an Olympus SZX microscope and transferred to soil.

### Histochemical localization of GUS and β-Gal staining

For X-gluc staining, X-GlcA ((5-bromo-4-chloro-3-indolyl-beta-D-glucuronic acid, Goldbio) in DMF (Dimethyl formamide)) was added to 50 ml of phosphate buffer (100 mM phosphate buffer saline with 100 mM Na_2_HPO_4_, NaH_2_PO_4_ each) and 50 mM K_4_FeCN_6_ and K_3_FeCN_6_ each, 0.5M EDTA and 10% Triton X-100. X-gluc was added to a final concentration of 100 mg/100 mL buffer. Harvested root tissue were vacuum infiltrated with the X-Gluc solution for 10 minutes and then incubated at 37 °C in dark for varying time periods. Two distinct GUS staining times were used for each construct at early nodulation time points (4 dpi, 10 dpi) and 28 dpi. These were *MtGLV1* (2 hrs, 2 hrs), *MtGLV2* (24 hrs, 24 hrs), *MtGLV6* (24 hrs, 12 hrs), *MtGLV9* (4 hrs, 4 hrs), *MtGLV10* (24 hrs, 4 hrs), *MtD14* (6 hrs, 6 hrs), *MtSPL5* (6 hrs, 24 hrs) and *MtMAKRL* (24 hrs, 6 hrs). At 28 dpi, nodules were excised and cleared with 1/20 strength bleach overnight and imaged.

Samples were washed in phosphate buffer three times before staining rhizobia with X-gal (Goldbio). Prior to X-gal (5-bromo-4-chloro-3-indolyl-β-D-galactopyranoside) staining, tissue was fixed in 2.5% glutaraldehyde by vacuum infiltration for ten minutes followed by a one-hour incubation time. Samples were rinsed with Z-buffer (100 mM Na2HPO4 and NaH2PO4 each, 10 mM Potassium chloride and 1 mM Magnesium chloride) and immediately transferred to the X-gal solution in Z-buffer (50mM each K4FeCN6 and K3FeCN6, 4% X-gal in DMF). Tissue was vacuum infiltrated, incubated in the dark overnight and imaged the next day.

### Hormone Treatments and Nodule excision

Peptides synthesis was carried out by Pepscan and Austin Chemicals, Inc. For auxin treatments, overnight germinated *M. truncatula* seedlings were transferred onto 1% water agarose media and grown for three days at 24°C. Sixty to seventy seedlings were transferred to sterile water (pH adjusted to 6.8) containing 1 uM Indole-3-acetic acid or equivalent amount of DMSO (Dimethoxy sulfoxide – solvent control) and treatment allowed to proceed for three hours. Post treatment for three hours, roots were excised from 20 seedlings per replicate per biological replicate and shoots discarded. For variable-N treatments, seedlings were grown for three days on B&D media with six mM nitrogen before transfer to liquid B&D medium at two different N-concentrations. Solution designated as Full-N constituted six mM nitrogen (Final concentration 0.5 mM KH_2_PO_4_, 0.25 mM K_2_SO_4_, 0.25 mM MgSO_4_, .01 mM Fe-citrate, 1 mM CaCl_2_, 2 mM KNO_3_, 2 mM NH_4_NO_3_, pH 6.8) while Low-N solution contained a limited amount of nitrogen, prepared without any NH_4_NO_3_ and only 0.5 mM KNO_3_.

### RNA Extraction, complimentary DNA synthesis and quantitative PCR

Total RNA was extracted using Trizol Reagent (Life Technologies) following the manufacturer’s recommendations (Invitrogen GmbH, Karlsruhe, Germany), digested with RNase free DNase1 (Ambion Inc., Houston, TX) and column purified with RNeasy MinElute CleanUp Kit (Qiagen). RNA was quantified using a Nanodrop Spectrophotometer ND-100 (NanoDrop Technologies, Wilington, DE). RNA integrity was assessed on an Agilent 2100 BioAnalyser and RNA 6000 Nano Chips (Agilient Technologies, Waldbronn, Germany). First-strand complementary DNA was synthesized by priming with oligo-dT_20_ (Qiagen, Hilden, Germany), using Super Script Reverse Transcriptase III (Invitrogen GmbH, Karlsruhe, Germany) following manufacturer’s recommendations. Primer Express V3.0 software was used for primer design. qPCR reactions were carried out in an QuantStudio7 (ThermoFisher Scientific Inc.). Five microliters reactions were performed in an optical 384-well plate containing 2.5 μL SYBR Green Power Master Mix reagent (Applied Biosystems), 15 ng cDNA and 200 nM of each gene-specific primer. Transcript levels were normalized using the geometric mean of two housekeeping genes, *MtUBI* (Medtr3g091400) and *MtPTB* (Medtr3g090960). Three biological replicates were included and displayed as relative expression values. Primer sequences are provided in **Supplemental Table 3**.

### Root Embedding and Sectioning

One cm root segments were fixed with 5 % glutaraldehyde in Phosphate Buffer Saline (pH = 7.2) solution overnight. Samples were rinsed three times and dehydrated using ethanol gradients (20 %, 40 %, 60 %, 80 % and 100 %). The samples were embedded in Technovit 7100 (Heraeus-Kulzer, Wehrheim, Germany), according to the manufacturer’s protocol. Samples were sectioned to 2.5 μm thickness using a microtome and stained with Toluidine blue for 1 min or till the desired color intensity developed and rinsed three times before imaging.

### Statistical Analysis

All statistical analyses were performed using GraphPad Prism 8 and tests selected therein. A two-sided Student’s t-test was used for comparison between genotypes or treatments. For multiple genotypes ordinary one-way analysis of variance tests or the Brown-Forsythe and Welch tests were performed followed by post-hoc statistical tests as mentioned.

### Figure Preparation and R packages used

Figures were prepared using Adobe Illustrator Creative cloud. Images were edited using Adobe Photoshop and FIJI. Graphs were prepared using GraphPad Prism 8. Phylogenetic tree was prepared using Mega X and the FigTree Application. All default plots were edited using Adobe Illustrator for clarity.

### Phylogenetic Tree Construction

Orthologs of GLV peptide encoding genes were retrieved from 18 different species by performing a Smith-Waterman search with SSearch(Ropelewski *et al*., 2003), and e-values of ≤ 0.01 were used for significant homologies followed by manual BLAST searches. Both the full protein as well as the short peptide coding region of known peptides in Arabidopsis and Medicago were used to initiate these searches. Retrieved sequences were selected through the SSP classification pipeline on MtSSPdb.noble.org. A maximum likelihood phylogenetic tree of the resulting list of putative GLV peptide coding proteins was generated using Mega X software and 1000 bootstrap iterations performed. Consensus tree was modified using Figtree and Adobe Illustrator.

## RESULTS

### Members of the GOLVEN peptide family are transcriptionally regulated by auxin, plant nitrogen-status, nodule organogenesis and the transcription factor NIN

By combining publicly available gene expression data from nodule segments at 4, 10, 21 and 28 days post inoculation (dpi) (de Bang *et al*., 2017; Boschiero *et al*., 2020) and quantitative reverse transcription PCR (qRT-PCR), we found that five of the fifteen members of the GOLVEN/ROOT MERISTEM GROWTH FACTOR (GLV/RGF) family, namely *MtGLV1, MtGLV2, MtGLV6, MtGLV9* and *MtGLV10* are more highly expressed in nodules than in roots at all nodulation stages analyzed (**Figure 1a, Supplemental Figure S1**). Three of these genes were induced by short term N-deprivation stress (0.5 mM NO_3-_) compared to 6 mM N (4 mM NO_3-_ + 2 mM NH_4+_) for 48 hours but not if the treatments were allowed to proceed longer (**Figure 1a, Supplemental Figure S1**).

**Figure 1.**
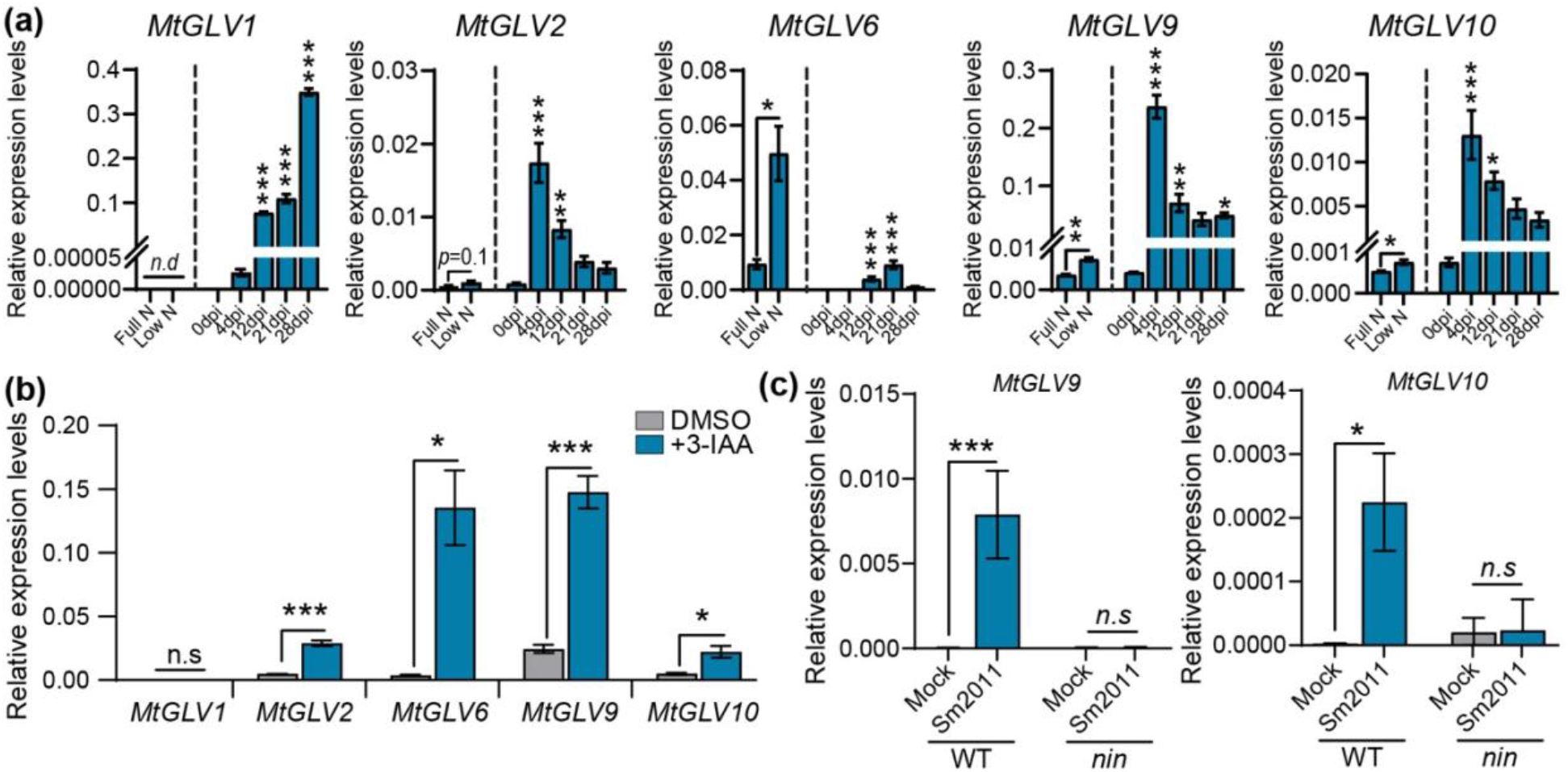
Expression of GOLVEN/ROOT GROWTH FACTOR peptide-coding genes is induced during nitrogen deficiency, auxin treatments and root nodule symbiosis. (a) Bar charts showing quantitative-PCR estimation of transcript abundance after 48 hours of nitrogen deprivation and at different time points (days post inoculation) during nodule development compared to uninfected roots (0 dpi). Error bars depict standard error of mean. (b) Bar charts showing quantitative-PCR estimation of transcript abundance in *M. truncatula* seedling roots treated with 1 μM auxin (3-IAA) or the solvent control (DMSO). Data are representative of three biological replicates with 40-60 seedling roots per replicate. Error bars depict standard error of mean. (c) Bar charts showing quantitative-PCR estimation of MtGLV9 and MtGLV10 transcript abundance in spot inoculated nodules on *M. truncatula* WT and *nin* mutants. Data are representative of three biological replicates each with at least 50-60 spot inoculated nodules.

Establishment of localized auxin maxima is required for lateral organ formation, and during nodulation is dependent on the transcription factor NODULE INCEPTION (NIN) (Leyser, 2018; Schiessl *et al*., 2019). Since GLVs act by altering auxin transport and signaling in Arabidopsis roots (Whitford *et al*., 2012; Fernandez *et al*., 2020), we tested whether their expression is regulated by auxin. Four of the five nodule-enhanced *GLVs* were upregulated in seedling roots treated with 1 μM Indole-3-acetic acid (3-IAA) three hours post treatment (**Figure 1b**). Further, expression data in MtSSPdb.noble.org show that *MtGLV9* and *MtGLV10*, but not other *GLVs* are induced in nodule primordia at 1 dpi. Using spot inoculated root segments infected with *Sinorhizobium meliloti* strain 2011 (Sm2011) for 24 hours, we detected that *MtGLV9* and *MtGLV10* transcripts were induced at nodule initiation sites in WT but not in *nin-1* mutants, indicating that induction of these genes is dependent on NIN (**Figure 1c**). These data suggest that expression of GLVs at nodule initiation sites occurs in a NIN dependent manner (Huo *et al*., 2006; Roy *et al*., 2017; Schiessl *et al*., 2019).

### Overexpression of five nodule induced *GOLVEN* genes negatively regulates nodule number

Spatial expression patterns obtained using promoter-GUS fusions in transgenic nodules corroborated the qRT-PCR data (**Figure 1**). The five *GLV* genes showed overlapping patterns of expression at sites of nodule initiation associated with successful infections and cell divisions suggesting they might act redundantly during nodule formation (**Figure 2**). Of the five genes, *MtGLV10* had the most confined expression pattern, being restricted to dividing cells underlying infection sites, while expression of *MtGLV1, MtGLV2* was associated with nodule vascular bundles closer to the nodule meristem (**Figure 2a, Supplemental Figure S1**). None of the five genes were expressed in infected root hairs (**Figure 2a, Supplemental Figure S1**). In mature nodules, expression of the GLVs was confined to the meristem, typical of genes involved in meristem maintenance as in Arabidopsis (Fernandez *et al*., 2015). We observed a similar but non-overlapping expression pattern of all five *GLV*s in initiating lateral roots (LRs) and the root tip. *MtGLV1* and *MtGLV10* were expressed in all dividing cells of the LR while *MtGLV2, MtGLV6, MtGLV9* were expressed on the flanks of LR primordia. At the root tip, *MtGLV9* and *MtGLV10* were expressed in columella cells, *MtGLV1, MtGLV2* were expressed in root cap cells and *MtGLV6* was expressed in the lateral root cap cells (**Figure 2a**).

**Figure 2.**
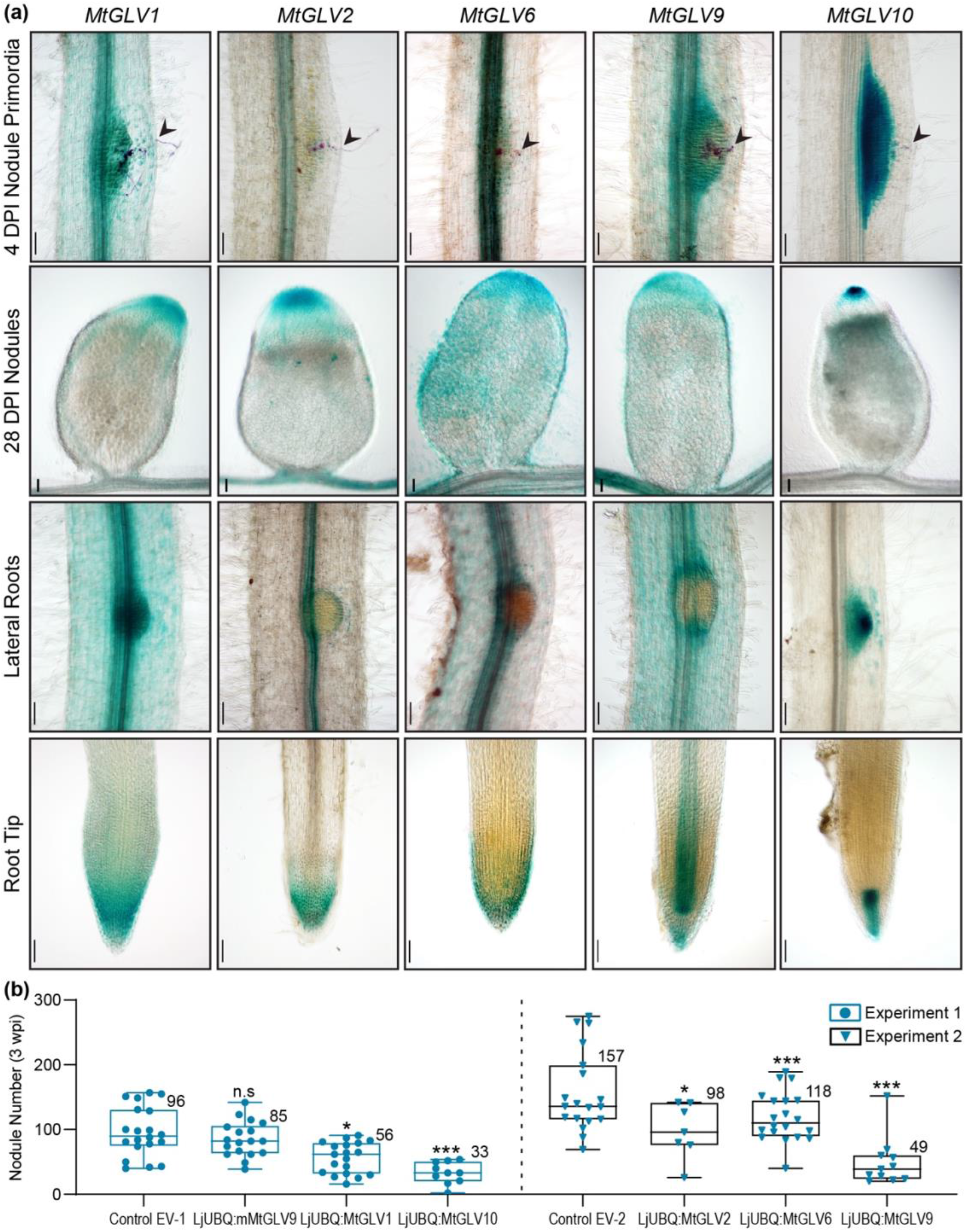
Five GOLVEN/ROOT GROWTH FACTOR peptide-coding genes are expressed during nodule organogenesis and root growth. (a) Promoter-GUS reporter fusions showing spatial expression of nodule induced *GLV* peptide-coding genes in nodule primordia, mature nodules, lateral roots and root tips. Arrowheads indicate infection threads. Scale bars denote 100 μM except for mature nodules (28 DPI) which measure 500 μM. At least 4-6 independent hairy root lines were assessed at every time point. (b) Overexpression of all five *GLV* coding genes in hairy roots of *M. truncatula* suppresses the formation of nodules. Each individual dot or triangle in the box plot indicates an independent line. Average nodule number of indicates on the shoulder of each box plot. One way ANOVA followed by Dunnett’s multiple comparison test was performed separately for experiment 1 and experiment 2 with their respective controls where \**p*<0.05, \*\**p*<0.01, \*\*\**p*<0.001. Data displayed summarize two independent experiments per construct. Average values are provided on the shoulder of each box plot.

To determine effects of GLV peptides on nodulation, we cloned the coding regions of *MtGLV1, MtGLV2, MtGLV6, MtGLV9* and *MtGLV10*, downstream of the Lotus *UBIQUITIN* promoter and generated transgenic hairy roots, via *Agrobacterium rhizogenes*-mediated transformation (Maekawa *et al*., 2008). We included a clone with a single base pair deletion in the coding region of *MtGLV9* (+203 bps from ATG) as a negative control along with an empty vector overexpressing the *GUS* gene. Consistent with peptide effects, overexpression of the *GLV* genes reduced the number of nodules formed on transformed roots 21 dpi by 25-50% (**Figure 2b, Supplemental Figure S4**). *MtGLV9* and *MtGLV10* had the strongest inhibitory effects on nodulation whereas the mutated *mMtGLV9* had no significant effect on nodulation.

### GOLVEN peptide encoding genes are present in genomes of all plants that form roots or root-like organs

To investigate neofunctionalization of these *GLV* genes in legumes, we retrieved their orthologues from twenty-one species of the Fabales, Fagales, Cucurbitales, Rosales nodule-forming eurosids, the rosid and asterid clades of the eudicots as well as monocots, gymnosperms and basal lycophytes, bryophytes and chlorophytes (**Figure 3, Supplemental Table S1**). All five nodulation-induced GLVs had putative orthologues in non-nodulating plant species suggesting no specialized roles during nodulation. However, although the *GLV* genes were present in plants that form roots or rhizoids, they were completely absent in the chlorophytes, which lack roots, such as *Chlamydomonas reinhardtii*, suggesting an evolutionary role in land colonization (**Figure 3)**, (Furumizu *et al*., 2021). In contrast with Furumizu et al., we did find putative GOLVEN orthologues in the lycophyte representative *Selaginella moellendorffii* genome, encoding putative thirteen amino acid long bioactive peptides (**Figure 3, Supplementary Table 1)**. The lycophyte clade is the most ancient clade with rooting plants.

**Figure 3.**
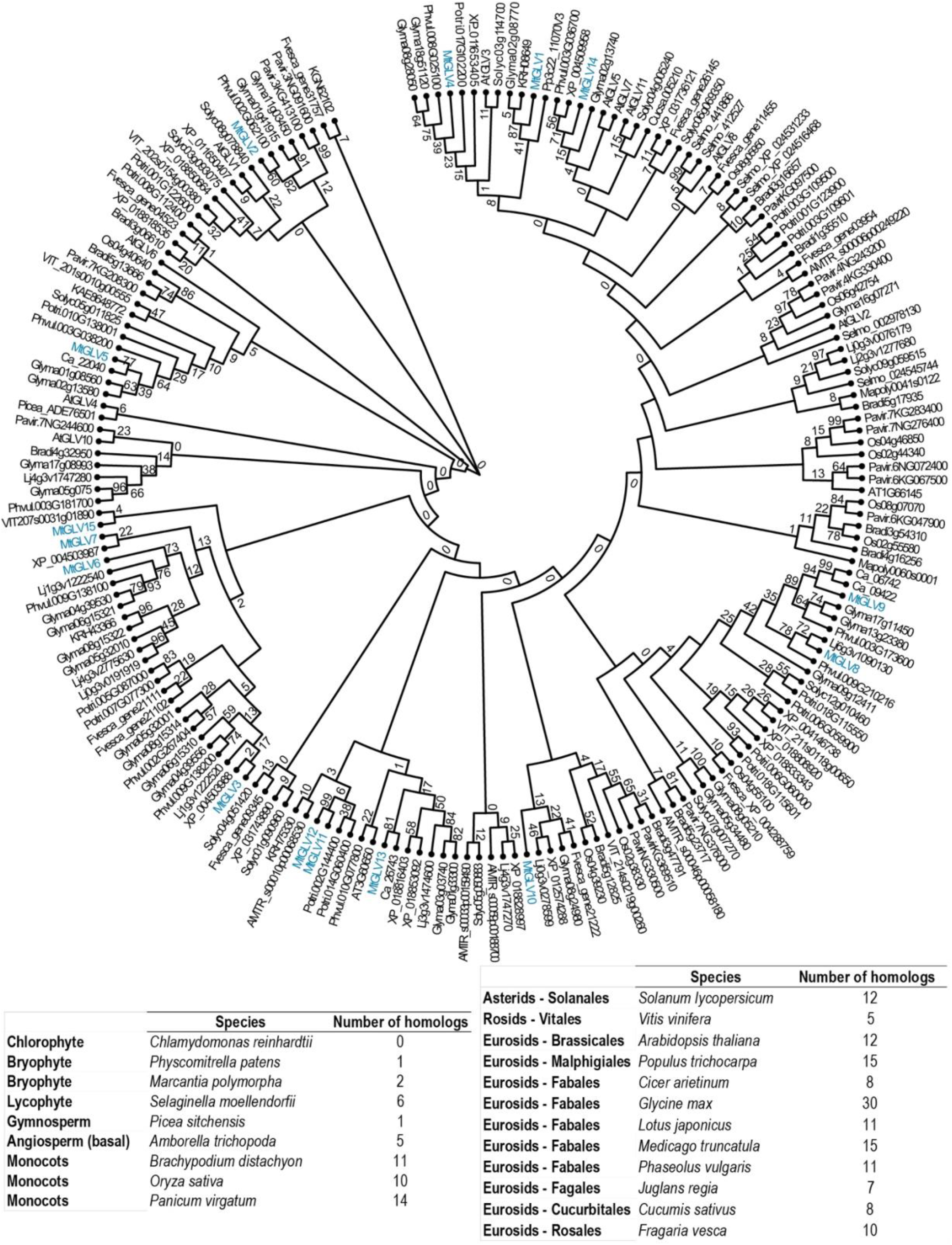
GOLVEN like peptides are encoded in all plants that form roots or root-like structures. Maximum likelihood phylogenetic tree created with the full length GOLVEN encoding polypeptide using MEGA X with 1000 bootstraps each. Please refer to Supplementary Table 1 for gene IDs and corresponding peptide sequence.

### Synthetic GOLVEN peptides regulate ten of eighteen root growth parameters tested

To determine if externally applied synthetic peptides phenocopied GLV function *in planta* we took a chemical genetics approach to explore the role of GLVs in root nodule symbiosis and control of root architecture traits. We synthesized peptide variants and measured their effects on root growth parameters and nodulation traits to identify the synthetic peptide with the strongest, most reproducible effects for further investigative studies. Peptides predicted to be encoded at the C-terminal domain of *MtGLV1, MtGLV2, MtGLV6, MtGLV9* and, *MtGLV10*, immediately following the peptidase cleavage recognition motif ‘DY’ were synthesized (**Supplemental Figure S3)**. Phenotypic effects of the peptides GLV1p, GLV2p, GLV6_hyp9p, GLV6_hyp10p, GLV9p, and GLV10p carrying a modified sulfotyrosine at position two, applied at 1 μM concentration, were distinct from those of scrambled, unmodified and unsulfated GLV10p, and solvent controls (**Figure 2A, B**). These results reinforce previous studies that show sulfation of the tyrosine at position two is essential for GOLVEN peptide activity (Matsuzaki, Y. *et al*., 2010).

GLV peptides significantly affected at least ten of the eighteen root growth parameters we tested (**Supplemental Table S2)**. As in Arabidopsis, GLV peptides had a positive effect on primary root growth/length (**Figure 2A)** and rendered the roots agravitropic to varying degrees with effects of GLV10p being the most visually distinctive (**Supplementary Figure S4, Figure 4a**) (Matsuzaki, Y. *et al*., 2010; Meng *et al*., 2012; Whitford *et al*., 2012). In contrast to their positive effects on root length, GLV1p, GLV2p, GLV6_hyp9p, GLV6_hyp10p, GLV9p, and GLV10p had negative effects on lateral organs, i.e., LRs and nodules, including organ number and density along the primary root (**Figure 4b, Supplemental Figure S3**). Clustering analysis of all traits revealed that GLV9p and GLV10p were the closest in terms of effects on root growth phenotypes (**Figure 4b**). Peptides did not affect *R. meliloti* Rm2011 growth rates over 60 hours indicating that nodulation phenotypes were due to effects on the plant and not its microsymbiont (**Supplemental Figure S3**). Interestingly, the position of the most basal nodule (closest to the shoot-root junction, developmentally oldest nodule, Trait 16) relative to the primary root length shifted upon GLVp treatment as nodules initiated more distally on the primary root (**Figure 4c**). However, because application of the GLV peptides reduced the total number of nodules formed. On the other hand, the more distal absolute position of the first formed nodule was highly consistent upon GLV10p application and resulted in a reduced absolute length of the zone over which nodules initiated (Trait 18) (**Figure 4c, Figure 5**).

**Figure 4.**
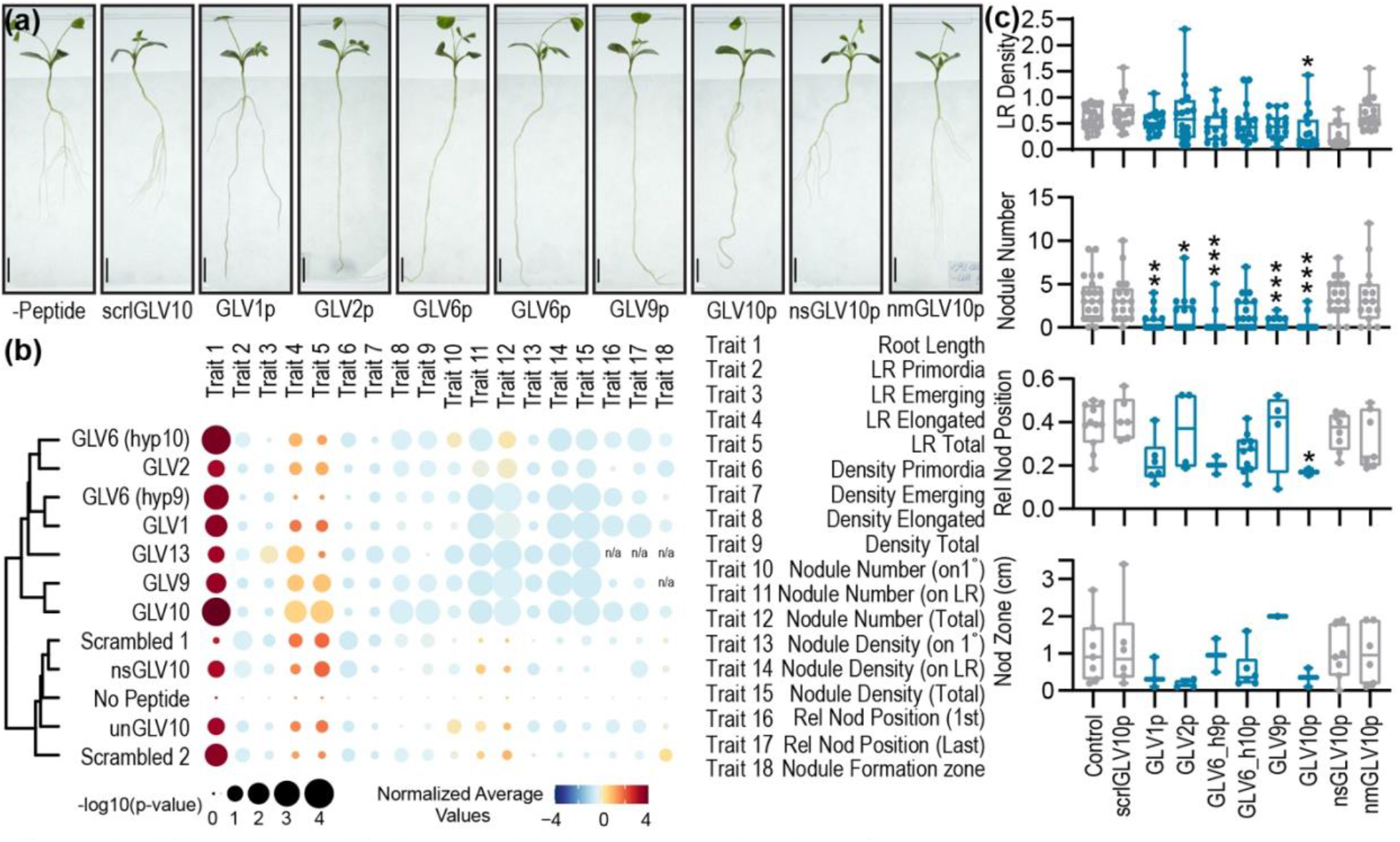
GLV peptides affect root architecture and nodule formation. (a) Representative images showing *M. truncatula* Jemalong A17 seedlings 10 days post germination (dpg) on plates containing 1 μM of the indicated peptide in B&D medium with 0.5 mM KNO3. The predicted bioactive peptides (GLVp) were around 13 amino acid residues long except for GLV2p (Supplementary Figure 2). Peptides were synthesized with a sulfotyrosine residue at position two and a hydroxyproline at position ten. In case of MtGLV6 where there was more than one proline, we designed alternate versions with a hydroxyproline in either position 9 (GLV10_hyp9) or 10 (GLV10_hyp10). A synthetic version of MtGLV13 derived peptide, which was not regulated during nodulation, was included. As negative controls we generated a non-modified (non-sulfated, non-hydroxylated) version of the most potent peptide GLV10p (nmGLV10) and a hydroxylated non-sulfated version (nsGLV10). We generated a ‘scrambled’ version of MtGLV10 (scrlGLV10) with identical amino acid composition but a randomized order of amino acids with or without the secondary group modifications (Refer to Extended Data Fig. 2 for corresponding peptide sequences). (b) Corresponding rain plot showing cumulative growth data of seedlings 10 dpi with *Sinorhizobium meliloti* Rm 2011 dsRED. Colors represent normalized average values with red representing increased trait values and blue representing decreased trait values. Data were compared using a Student’s t-test and bubble size indicates -log(p-value). Refer to Supplementary Table 1 for trait definitions. (c) Boxplots of four individual traits of interest showing effect of GLV peptides compared to negative controls; Trait 9 (Total LR density), Trait 12 (Total nodule number), Trait 16 (Relative position of first formed nodule), Trait 18 (Nodule formation zone). Asterisks represent *p<0.05, **p<0.01, ***p<0.001 using a Brown-Forsythe and Welch ANOVA protected Dunnett’s multiple comparison test (treatment vs no-peptide control).

**Figure 5.**
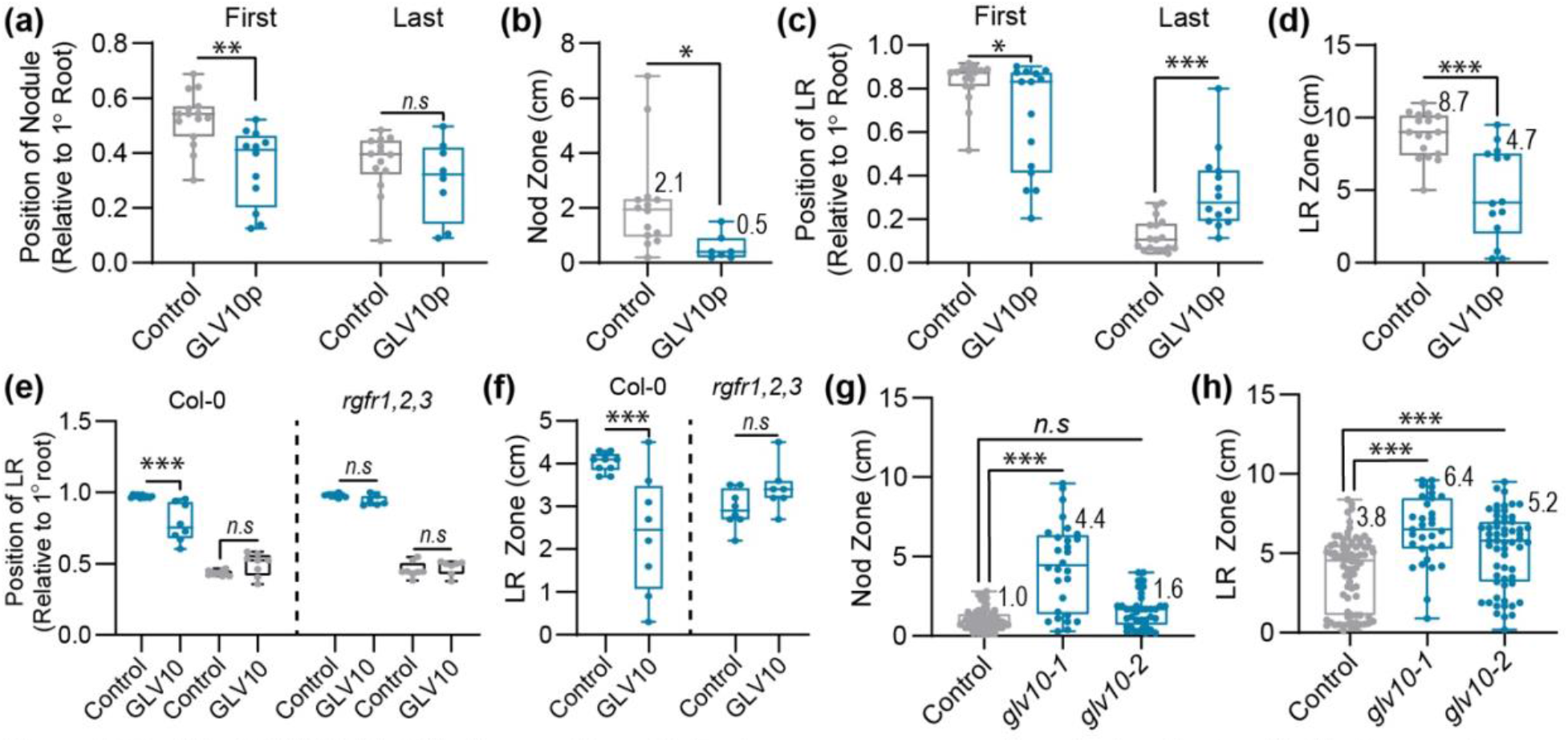
Peptide GOLVEN10 shifts the position of lateral organs consequently reducing the longitudinal zone of organ formation. Position of the developmentally first and last formed organ relative to the primary root length measured from the root tip with and without GLV10p treatment and the resulting zone of organ formation on *M. truncatula* Jemalong A17 seedlings 14 dpi with Sm2011 dsRED and 17 days post germination. (a) Nodule position (b) Nodule formation zone (c) Lateral root position (d) LR formation zone. Asterisks represent \**p*<0.05, \*\**p*<0.01, \*\*\**p*<0.001 using a Student’s t-test. (e,f) Position of the developmentally first and last formed organ measured from the root tip, relative to the primary root length with and without 10 μM GLV10 peptide treatment and the resulting zone of organ formation on *A. thaliana* WT Col-0 and *rgfr1,2,3* mutant lines 14 days post germination. Asterisks represent \**p*<0.05, \*\**p*<0.01, \*\*\**p*<0.001 using ANOVA-protected Tukey’s multiple comparison test. (g,h) Size of the Nod Zone and LR Zone in WT R108 compared to single glv10 mutants two wpi with Rm2011. Data are representative of cumulative values from two identical experiments. Asterisks represent \**p*<0.05, \*\**p*<0.01, \*\*\**p*<0.001 using ANOVA-protected Dunnett’s multiple comparison test. Numeric values for the absolute zone of lateral organ formation are provided on the shoulder of box plots.

Since nodulation and rhizobial infections are genetically distinct processes, we tested effects of the peptides on early infection structure development. GLV peptides reduced the total root length, including secondary and tertiary LRs, as well as the total number of early infection events (**Supplemental Figure S2**). However, the GLVs did not affect early infection initiation events per cm root, except for GLV10p, which resulted in more microcolonies and infection thread initiations, but a strong reduction in nodule formation (**Supplemental Figure S2**). Increased infection induced by GLV10p may have been a consequence of the severe reduction of nodule number, as observed for symbiotic mutants with colonization defects such as *nf-ya1* (Laporte *et al*., 2014). As GLV peptides did not seem to affect infection directly but reduced nodule density, we focused on their effects on root cortical processes such as organogenesis.

### GLV10 controls the zone of lateral organ formation in an RGFR receptor-dependent manner

Lateral root positioning (rhizotaxis) is an understudied trait while nodule positioning over the longitudinal root axis (we term nodulotaxis) has not been described in literature before. Therefore, we further investigated these two traits. In Arabidopsis, periodic pulses of auxin-induced gene expression occur over a zone of the root close to the root tip called the ‘oscillation zone’ which pre-patterns longitudinal spacing or LR primordia positioning and consequently determines the zone of the root that is primed or capable of initiating lateral roots (Hofhuis *et al*., 2013; Laskowski & ten Tusscher, 2017). To better understand the effect of GLV peptides on lateral organ positioning (Traits 16-18), we pursued the peptide with the strongest and most reproducible effect, i.e., GLV10p (**Figure 4c**). In untreated seedlings, nodules typically originated at the midpoint (50%) relative to the length of the primary root. Upon GLV10p treatment, the position of the first nodule shifted to approximately 60% of the primary root length (**Figure 5 a,c**). Similarly, the relative position of the first lateral root was lower on the primary root, while that of the developmentally last formed root primordia appeared to be proximal to the root base (**Figure 5a**). This shift in organ positioning resulted in a statistically significant reduction of the absolute length of the root over which nodules formed (nod zone) and the zone that supported lateral roots (LR Zone, **Figure 5 b,d**). In cases where there were only two nodules on the main root, the distance between two successive nodules was reduced. We tested the effect of GLV10p peptide effects over the 1nM to 10 μM range on WT *M. truncatula* seedlings (**Supplemental Figure S5**) and found that effects were detectable even at a concentration of 100 nM upto 1 nM, depending on the trait under study.

In *A. thaliana*, GLV peptides are perceived by five LRR-RLKs *ROOT GROWTH FACTOR1 INSENSITIVE/ROOT GROWTH FACTOR RECEPTOR*, namely AtRGI1/AtRGFR1, AtRGI2/AtRGFR2, AtRGI3/AtRGFR3, AtRGI4, and AtRGI5. The Arabidopsis *rgfr1 rgfr2 rgfr3* triple mutant has a stunted primary root and an enlarged root meristem but retains sensitivity to GLV peptide with respect to LR density (Shinohara *et al*., 2016). We tested the effect of Medicago GLV10p on Arabidopsis root growth. GLV10p, which has three amino acids different from AtGLV10p, was perceived by Arabidopsis roots and, as in Medicago, shifted the relative position of the first lateral root to a more distal position on the primary root and decreased the zone of LR formation (**Fig 5 e,f**). However, the triple *rgfr* mutant was insensitive to GLV10p peptide with respect to LR positioning and LR zone (**Figure 5 g**), indicating that the effect of the peptide on LR zone formation was dependent on the RGI receptors.

Finally, to test whether these effects were also mediated by GLV10 *in planta* we isolated homozygous mutants of *MtGLV10, glv10-1* (NF12742) and *glv10-2* (NF20983), with exonic insertions of the *Tnt1* retrotransposon (**Supplemental Figure S3**) (Tadege *et al*., 2008). The lateral root zone appeared to be wider than the WT for *glv10-1* and *glv10-2*, but varied between alleles for the Nod zone (**Figure 5 g,h**). Since there are several other GLV encoding genes involved in nodulation, higher order mutants are necessary for characterizing these traits in depth. Taken together, these data indicate that GLV peptides are negative regulators of lateral root zone formation and that this may be dependent on RGFR receptors.

### Increased primary root growth upon GLV10 application is mediated by an increase in cell number per cortical cell file but not cell length

The spatial distribution of LRs or rhizotaxis is determined by a combination of three factors - primary root growth, organ density and the distance between successive organs (Du & Scheres, 2018). Studies in Arabidopsis show that application of AtGLV6 peptide to roots disrupts early symmetrical cell divisions required for correct lateral root initiation (Fernandez *et al*., 2020). Application of the peptide GLV10 increases root length and decreases lateral root density along the primary root both, in Arabidopsis and Medicago (**Figure 4b,c**). GOLVENs therefore control at least two of the three factors responsible for spatiotemporal organ positioning. To understand the cellular basis of GLV10 mediated root elongation in Medicago we collected longitudinal sections of roots 1 cm above the root tip which revealed that both cell number and cell size were affected by application of GLV10. While cell length along the longitudinal axis decreased by more than 50%, the cell number per cortical cell file increased by 50% (**Figure 6b**); this effect persisted even in mature regions of the root (**Figure S6**). Multicellular processes such as root growth are dependent on the combined activity of two linked processes, cell expansion and cell division (Jones *et al*., 2019). Faster root elongation rates are correlated with increased cell division at the root meristem, with little change in cellular expansion rates (Beemster & Baskin, 1998). In keeping with this observation, we propose that an increase in cortical cell number caused by GLV10 plays a role in regulating root tissue growth and consequently lateral organ positioning (**Figure 6c**).

**Figure 6.**
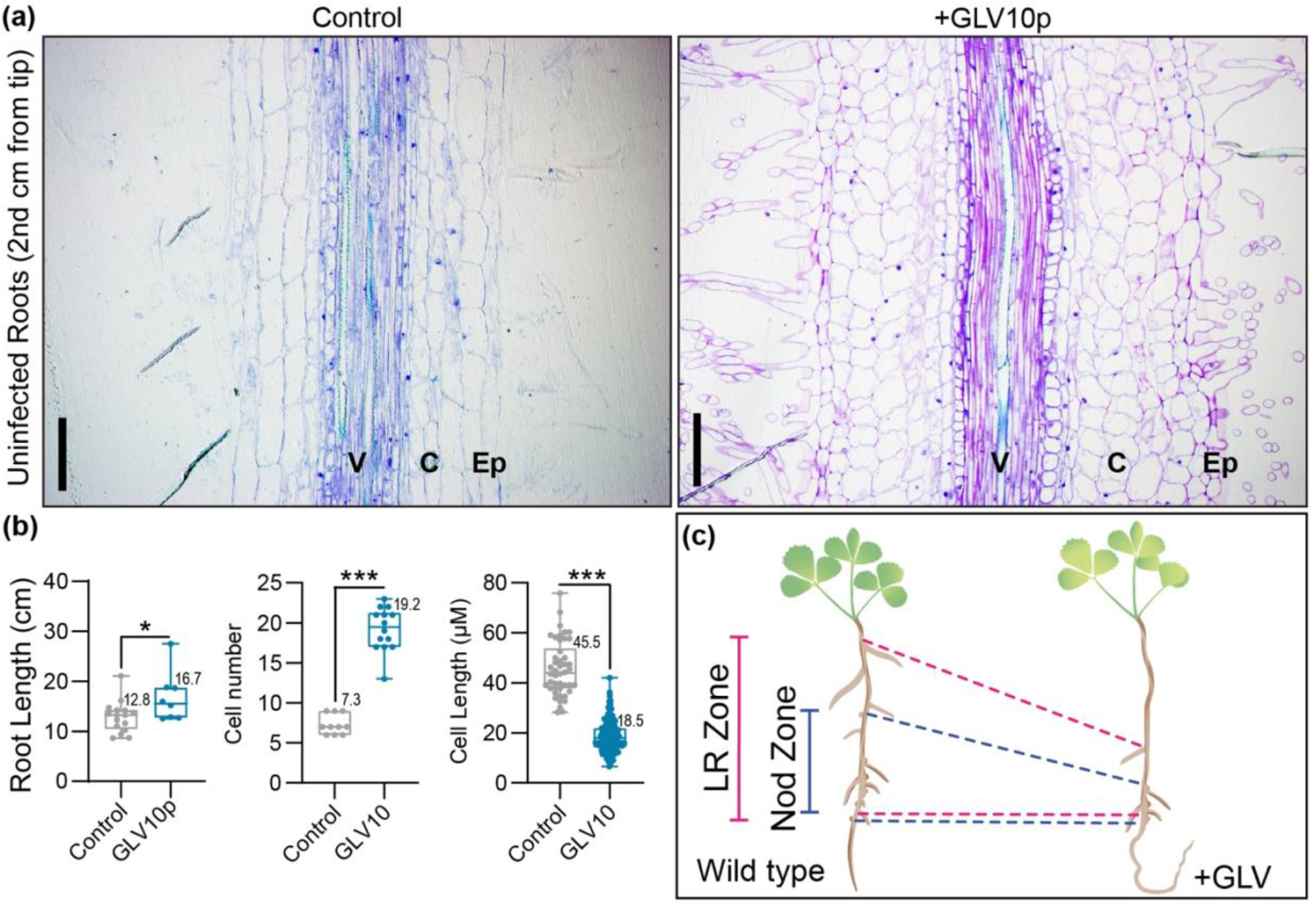
The synthetic peptide GLV10 affects cell number and cell size in Medicago roots. (a) Images show 2.5 μm thick sections of root segments collected one cm below root tip at a 20x magnification. Control roots treated with solvent (left) and roots treated with 1 μM GLV10 peptide (right) for seven days. Scale bars represent 50 μm. Segments from at least eight roots per sample were analyzed. (b) Quantification of data shown in (a) using ImageJ. Application of GLV10 peptide increases root length by increasing cell number but decreasing cell size in each cortical cell file. Roots were treated with peptide for seven days on plates and compared to untreated roots. Student’s t-test ***p<0.001. (c) Diagrammatic representation of nodulotaxy as mediated by the peptide GLV10.

## DISCUSSION

When nitrogen availability is low, plants require an internal signal(s) to inhibit LR production and promote primary root elongation in search of deeper, N-rich soil layers. *MtGLV10* and *MtGLV9* are peptide hormones induced in roots upon short term N-starvation that act as negative regulators of LR initiation and positive regulators of root elongation (**Figure 1a, Figure 4b**,**c**) phenocopying the N-starvation root architecture response (Mohd-Radzman *et al*., 2013). Modulation of root system architecture by GLVs in conjunction with auxin (**Figure 1b**) may have been instrumental in colonization of land by plants, allowing them to integrate nutrient stress signals into root developmental programs, given that GLVs are conserved in all land plants but not in rootless chlorophytes (**Figure 3**). Our data suggest that at least five GOLVEN peptide coding genes act in concert to control root nodule symbiosis and root architecture traits (**Figure 1, 2**). Of these five, MtGLV10 acts not only at the lateral organ initiation stage but earlier, likely during priming and organ positioning, which determines the zone of the root over which lateral organs can initiate and then emerge (**Figure 4, 5**). We apply the definition of the ‘Nodulation Zone’, as defined by Yoshida et al., as the zone along the longitudinal root axis that is capable of undergoing cell division and supporting formation of nodules (Yoshida *et al*., 2010). Similarly, the Lateral Root Zone, as interpreted in this study describes the zone along the longitudinal root axis that is capable of undergoing cell division leading to initiation of lateral roots. Like rhizotaxis, a phenomenon which ensures LR positioning and regular spacing between successive LRs, we find that a similar mechanism exists for nodule spacing, which we call nodulotaxis (**Figure 4a, b**). Although nodules can cluster together at a particular infection site, they are typically spaced out along the longitudinal root axis or the nodulation zone. Such spacing is lost, however, in the *sickle* mutant that forms a continuous chain of nodule primordia, many of which fail to develop into functional nodules (Penmetsa & Cook, 1997; Penmetsa *et al*., 2003). Thus, mechanisms controlling nodule positioning and priming are important to ensure formation of fully-developed, functional nodules. At present, rhizobial infection, nodule number and N-fixation efficiency are the predominant phenotypic traits that are measured when trying to understand gene function during root nodule symbiosis. Nodulotaxis as a trait is rarely considered, if not completely overlooked, but is nevertheless important to restrict nodule numbers to levels that can be supported effectively by plant photosynthesis. Our study identifies GLV10 as a dual regulator of both rhizotaxis and nodulotaxis that controls the zone of lateral organ formation in *M. truncatula*. Shifting the position of the first lateral organ more distal to the root base requires deferred organ initiation in space and/or in time. From studies in Arabidopsis, we know that GLV peptides disrupt initial cell divisions in the root pericycle, which delays or halts lateral root initiation (Fernandez *et al*., 2015; Fernandez *et al*., 2020). We found that in *M. truncatula*, another factor that contributes to this delay is faster root growth upon GLV application caused by an increase in cell number along the longitudinal root axis **(Figure 6b)**. Both Nodules and LRs initiate acropetally, and no new organs develop between already developed lateral organs (Dubrovsky *et al*., 2006). In theory, if treated roots are longer than those of control plants at the time of organ initiation, which only occurs in a narrow zone of the root close to the Root Apical Meristem, the first formed lateral organ will initiate more distally from the shoot compared to untreated controls. Rapid cell division stimulated by GLV10 accelerates root growth rates thereby spatially shifting the position of the first lateral organ formed **(Figure 5)**. Since organ positioning is determined by a combination of three factors i.e primary root length, organ density and the distance between successive organs (Du & Scheres, 2018), further temporal delay in initiation of subsequent lateral organs would facilitate an overall decrease in the LR zone length or Nod zone length caused by altered cell division at organ initiation sites (Fernandez *et al*., 2015). Fixed or flexible threshold theory for cell division which proposes that cells undergo division only after they have reached a certain threshold of size, would suggest that smaller cells in GLV10 treated roots may be unable to undergo asymmetric cell divisions until they have expanded sufficiently, impairing lateral organ initiation (Jones *et al*., 2019).

Low-N (<1 mM N) availability promotes the establishment of root nodule symbiosis, while simultaneously inducing the expression of *GLV*s that, ultimately, limit excessive nodulation (**Figure 1a**). Both, low N in roots and nodule development lead to GLV production in these organs similar to known Autoregulation of Nodulation pathway where CLE-SUNN signaling ensures sufficient but not excessive nodulation (Okamoto *et al*., 2013; Nishida *et al*., 2018). MtNIN directly induces the expression of *MtCLE13*, and mobile MtCLE13 is recognized by MtSUNN in the shoot, which mediates systemic AON that limits further nodulation (Laffont *et al*., 2020). Our data indicate that in addition to controlling the CLE (and CEP) nodulation regulators (Laffont *et al*., 2020), NIN is required for *MtGLV9 and MtGLV10* expression at nodule initiation sites (**Figure 1c**). Identification of GLV receptor(s) will enable investigations into possible crosstalk with the CLE-SUNN module that regulates nodule number through long distance signaling, or other root components such as SKL that control nodule number and positioning locally. Given that the *A. thaliana rgfr* triple mutant is insensitive to high doses of GLV10 peptide, which normally reduces the LR zone and alters LR positioning (**Figure 5 e,f**), the corresponding receptor orthologues in *M. truncatula* are interesting candidates for further investigations.

A question that remains unanswered is whether the nodulation zone is identical to the rhizobial infection susceptible zone given that nodule organogenesis and rhizobial infection are genetically separable processes. The finding that GLV10 application reduces nodule density over the total root length without changing infection thread density (**Figure S2**), combined with the observation that none of the five GLVs are expressed *in vivo* within infected root hairs (**Figure 2, Figure S2**) suggests that GLV10 controls cortical processes that regulate organogenesis rather than infection. Nevertheless, since the zone of infection competent root hair cells is narrower compared to the total root length supporting root hairs, further studies are required to understand GLV control of rhizobial infection including changes in root hair length, root hair density and length of the susceptibility zone upon GLV10 application.

In conclusion, our study introduces a new set of players, namely the GOLVEN signaling peptides, into the story of root nodule symbiosis. Our work reinforces chemical genomics as a powerful approach to understanding peptide function in plant development, such as nodulotaxis. Similar synthetic tools continue to be instrumental in understanding function of classical phytohormones in hormone research. Although, a lot remains to be discovered, our work presented here sets the stage for future work on this interesting new family of nodulation regulators.

## ACKNOWLEDGEMENTS

Authors gratefully acknowledge the help of Lynne Jacobs in the greenhouse, Lloyd Noble summer scholar Miss Sarah Dysinger for data entry and assistance with plant growth. We thank Dr. Julia Frugoli for the gift of the *sunn* mutant seeds and Dr. Yoshikatsu Matsubayashi for the gift of *rgfr1,2,3* mutant seeds. This research was funded by of NSF#1444549 and the Noble Research Institute.

## AUTHOR CONTRIBUTIONS

S.R, I.T.J, S.Z, W.L, K.S, H.K.L, C.B performed experiments and helped acquire data, S.R analyzed data, S.R, W.R.S, M.U conceptualized this study, P.X.Z, G.E.D.O, J.D.M, W.R.S, M.U supervised the research, S.R, M.U wrote the manuscript.

## DATA AVAILABILITY

All peptide gene overexpression constructs have been deposited with Addgene and can be accessed under deposit ID 79021.

## SUPPLEMENTAL INFORMATION

**Supplemental Table S1:** List of orthologous GOLVEN peptide coding genes found across 21 plant species.

**Supplemental Table S2:** Root architecture and nodulation described in this study.

**Supplemental Table S3:** List of Primers, constructs, and lines used in this study.

## SUPPLEMENTAL FIGURES

**Figure S1.**
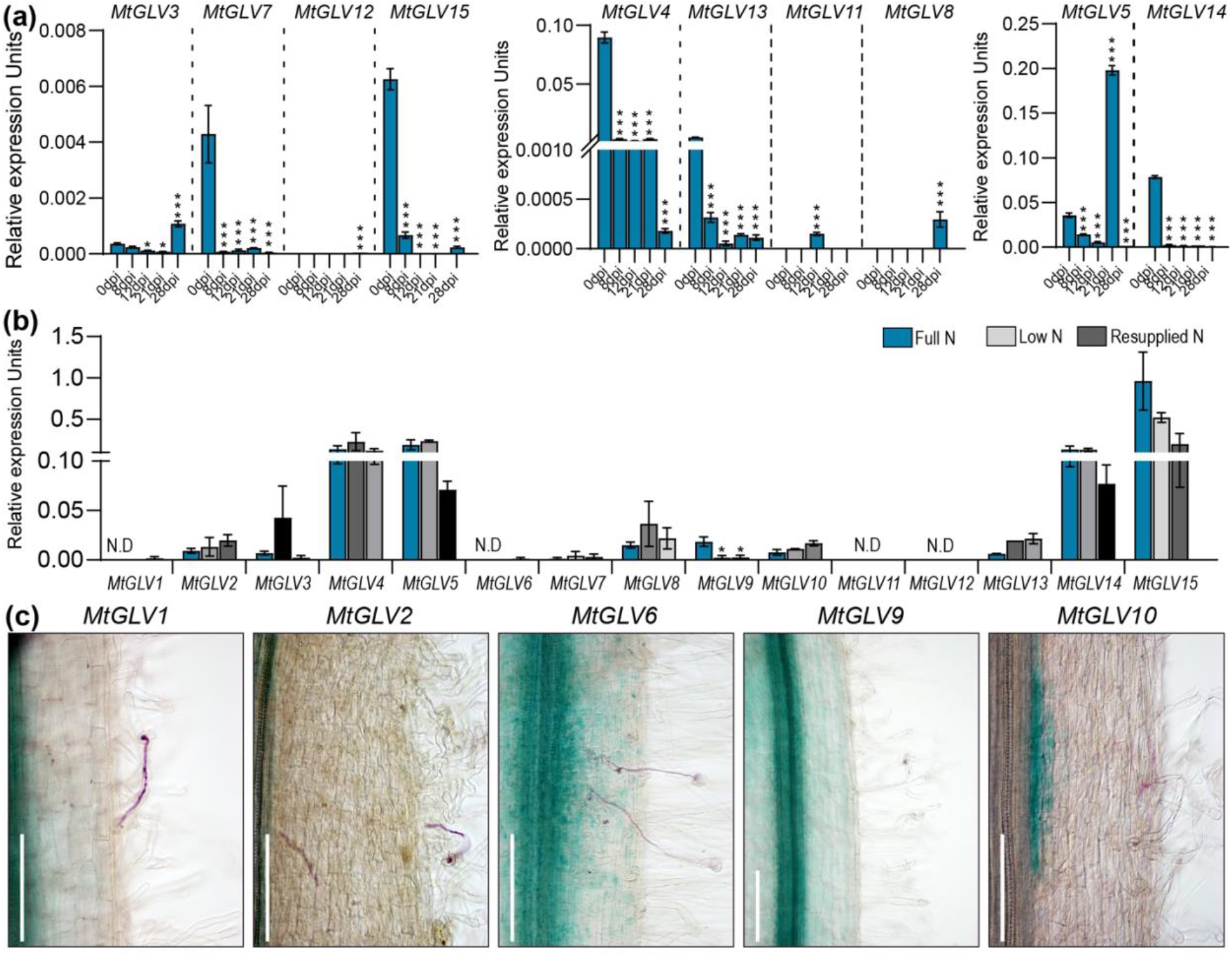
Expression of GLVs during nodulation, N-deprivation and in infected root hairs. Expression of GOLVEN peptide encoding genes in Medicago. (a) Quantitative PCR estimation of GLV transcript abundance at the denoted timepoints. *p<0.05, ***p<0.001 based on an ANOVA-protected Dunnett’s multiple comparison test (vs 0 dpi uninfected roots). Error bars indicate SEM and three biological replicates per time point were used. (b) qPCR estimation of GLV transcript abundance in *M. truncatula* A17 plants deprived of N for two weeks compared to plants supplemented with potassium nitrate as in de Bang et al., 20178. Student’s t-test *p<0.05. Error bars indicate SEM and three biological replicates per treatment were used. (c) GLV promoter-GUS reporter activity is absent in infection threads of M. truncatula hairy roots transformed with the indicated constructs four dpi with Rm1021. Rhizobia are co-stained in magenta-gal. Scale bars represent 100 μm.

**Figure S2.**
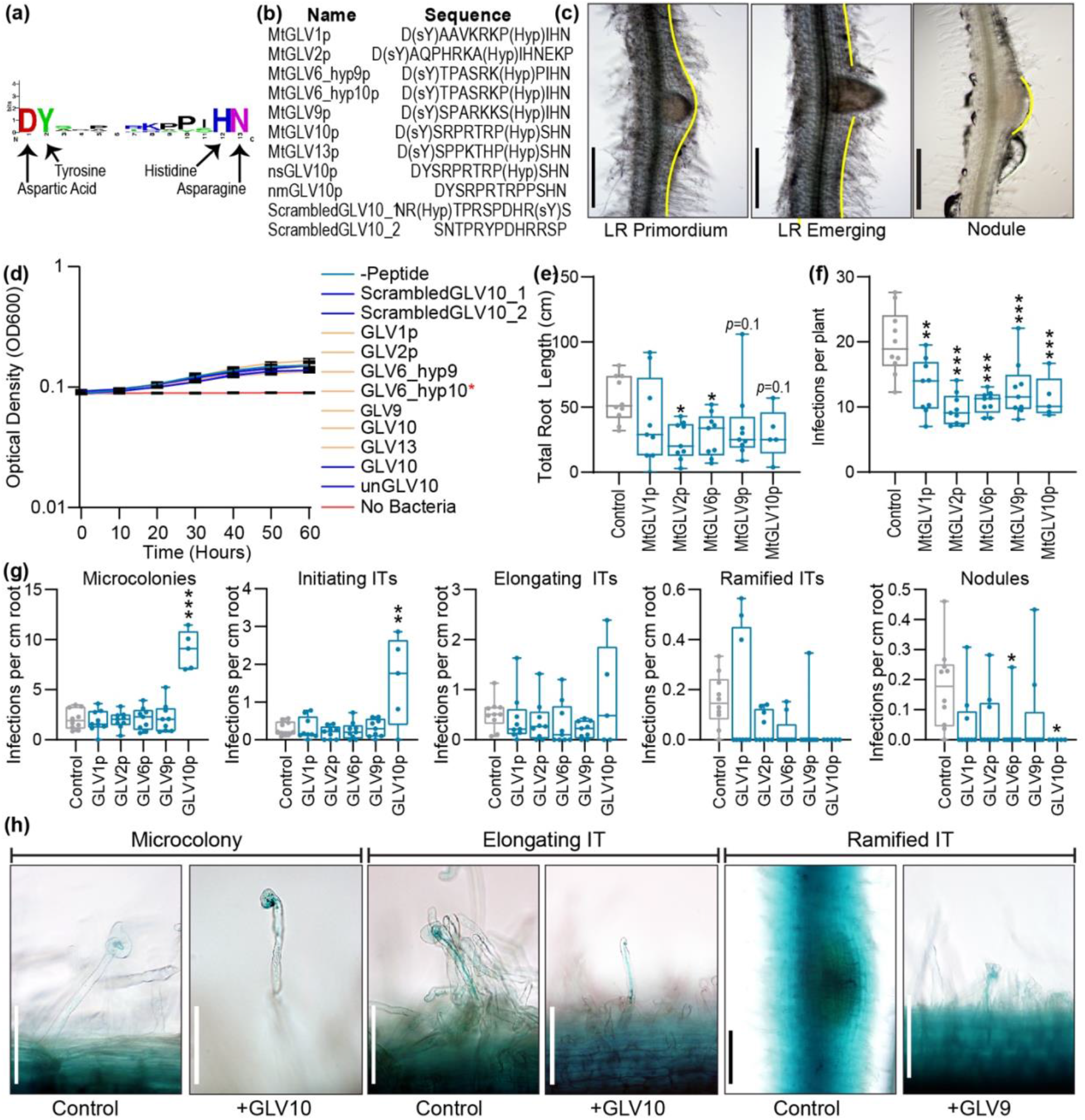
Sequence and physiological effects of synthetic GLV peptides used in this study. (a) Logo showing conserved residues in Medicago peptides. (b) Sequence of GLV peptides synthesized in this study. (c) *M. truncatula* root images showing stages of lateral root or nodules scored in this study. Scale bars denote 500 μm. See Supplementary Table 1 for trait definitions. (d) Time plots over 60 hours showing effects of synthetic peptides used in this study on growth of Rm2011 dsRED in the presence of the peptides as indicated. Asterisk * indicates a significant difference for GLV6_hyp10 which was not reproducible in subsequent experiments. (e) Change in total root length in *M. truncatula* Jemalong A17 seedling roots upon peptide treatment compared to control (no peptide). Asterisks represent \**p*<0.05 using a posthoc Dunnett’s multiple comparison test following a one-way ANOVA. (f) Change in total rhizobial infections upon peptide treatment compared to control in the same experiment. Asterisks represent \*\**p*<0.01, \*\**p*<0.001 using a posthoc Dunnett’s multiple comparison test following a one-way ANOVA. (g)Number of infection events per cm total root in the same experiment as (e,f) above. Asterisks represent \*\**p*<0.01, \*\**p*<0.001 using a ANOVA-protected Dunnett’s multiple comparison test. (h) Images showing infection structures in *M. truncatula* seedlings infected with Rm2011 HemA::LacZ seven days post inoculation. Scale bars represent 100 μm.

**Figure S3.**
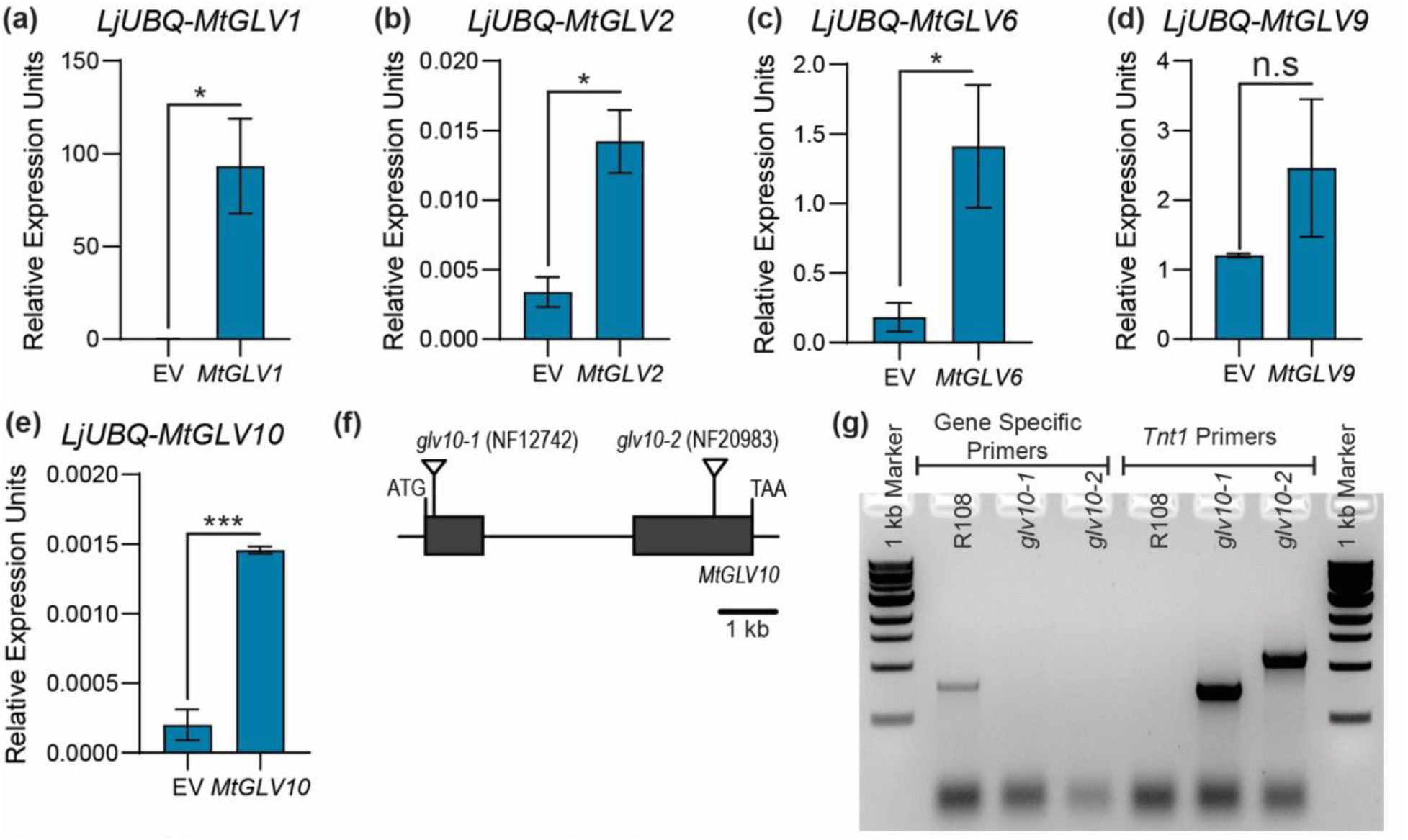
Characterization of lines used in this study. Expression of individual MtGLV genes in their corresponding over expression lines. (a) *MtGLV1* (b) *MtGLV2* (c) *MtGLV6* (d) *MtGLV9* (e) *MtGLV10*. Data represent qPCR estimation of transcript abundance using hairy root tissues. Error bars indicated SEM, n=2-4 per line. Student’s t-test \**p*<0.05, \*\*\**p*<0.001. (f) Gene structure showing position of *Tnt1* insertions in exonic regions of *glv10* mutant lines used in this study. (g) Agarose gel images showing PCR amplicons in WT R108 compared to mutants.

**Figure S4.**
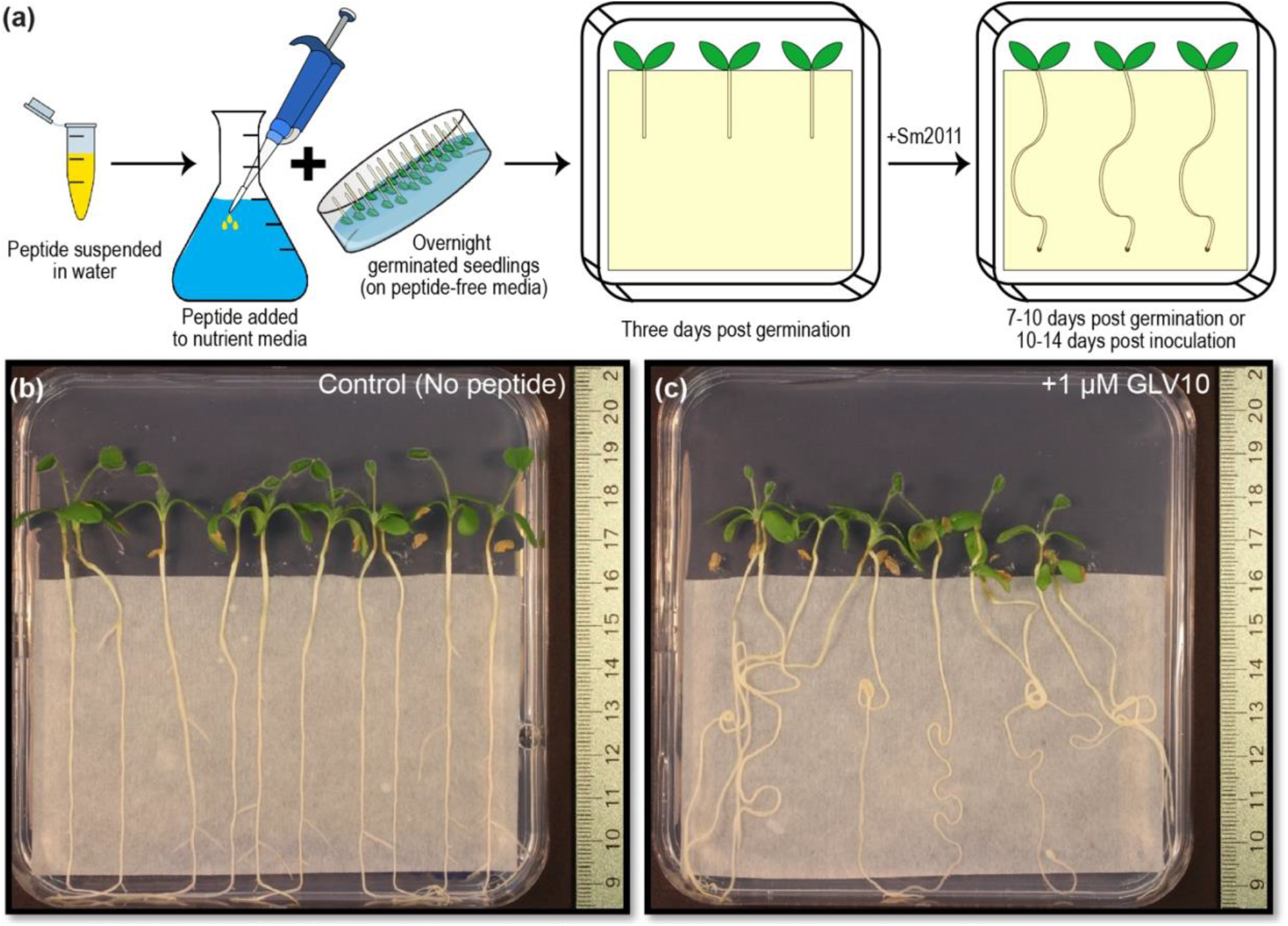
Effect of GLV10 peptide application on *M. truncatula* root growth. (a) Overview of peptide treatment and plant growth setup used in this study. (b, c) Representative images comparing effects of GLV10 peptide application to roots at 1 μM concentration compared to untreated roots. Images were taken 10 days post transfer to plates containing 1% Agarose in water.

**Figure S5.**
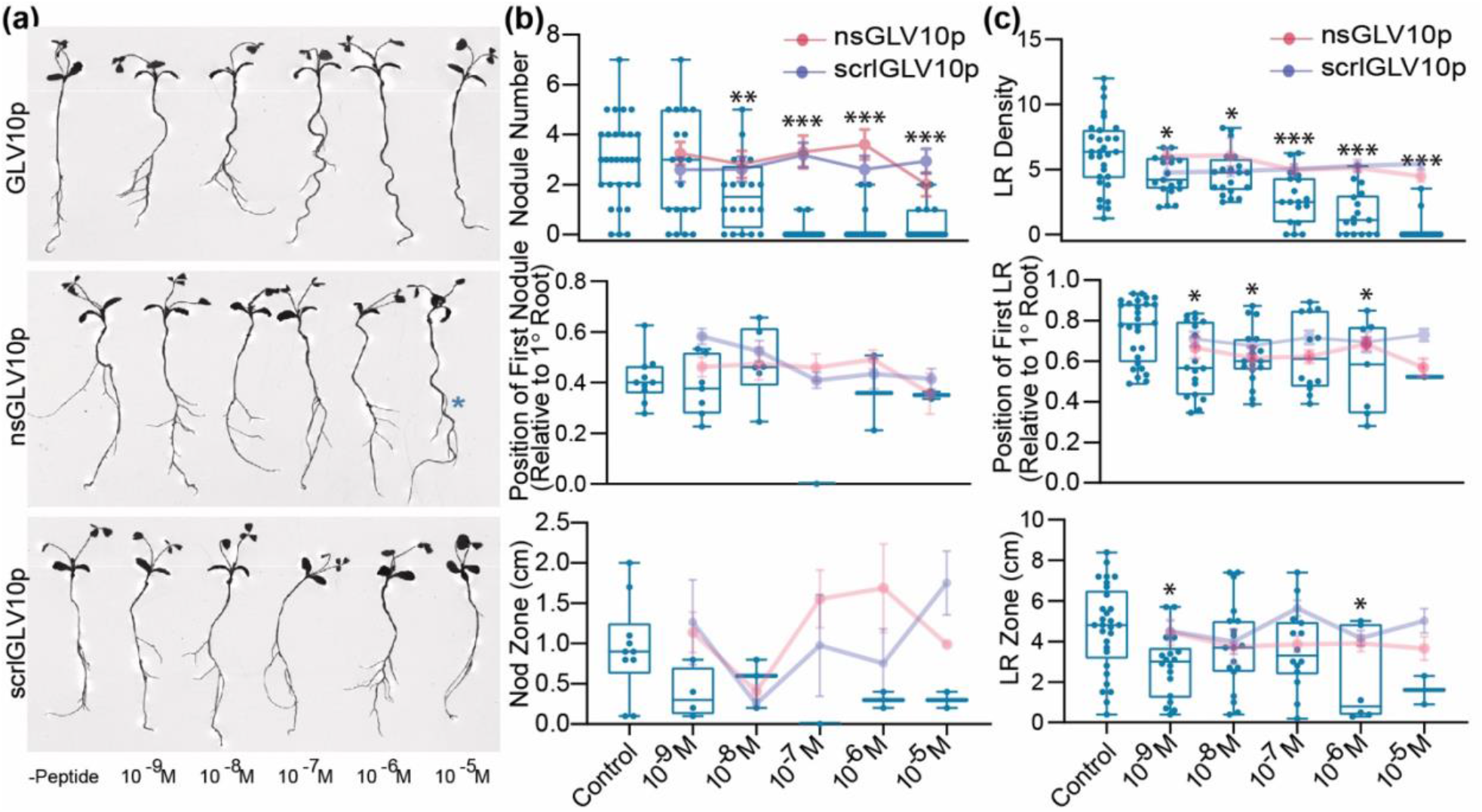
Peptide dilution curve. (a) Representative root scans showing seedling morphology 13 days post growth on B & D low-N media containing GLV10p, a modified non-sulfated version of the same peptide (nsGLV10p) and a scrambled version of the peptide (scrlGLV10p) at the concentrations indicated. See Supplementary Figure S3 for sequence details. (b) Number of nodules, position of developmentally first formed nodule relative to primary root length and the resulting nodule formation zone 10 days post inoculation with *S. meliloti* strain Rm2011 dsRed at the concentrations of peptides indicated. (c) Density of lateral roots formed, position of developmentally first formed lateral root relative to primary root length and the resulting LR formation zone in the same experiment.

**Figure S6.**
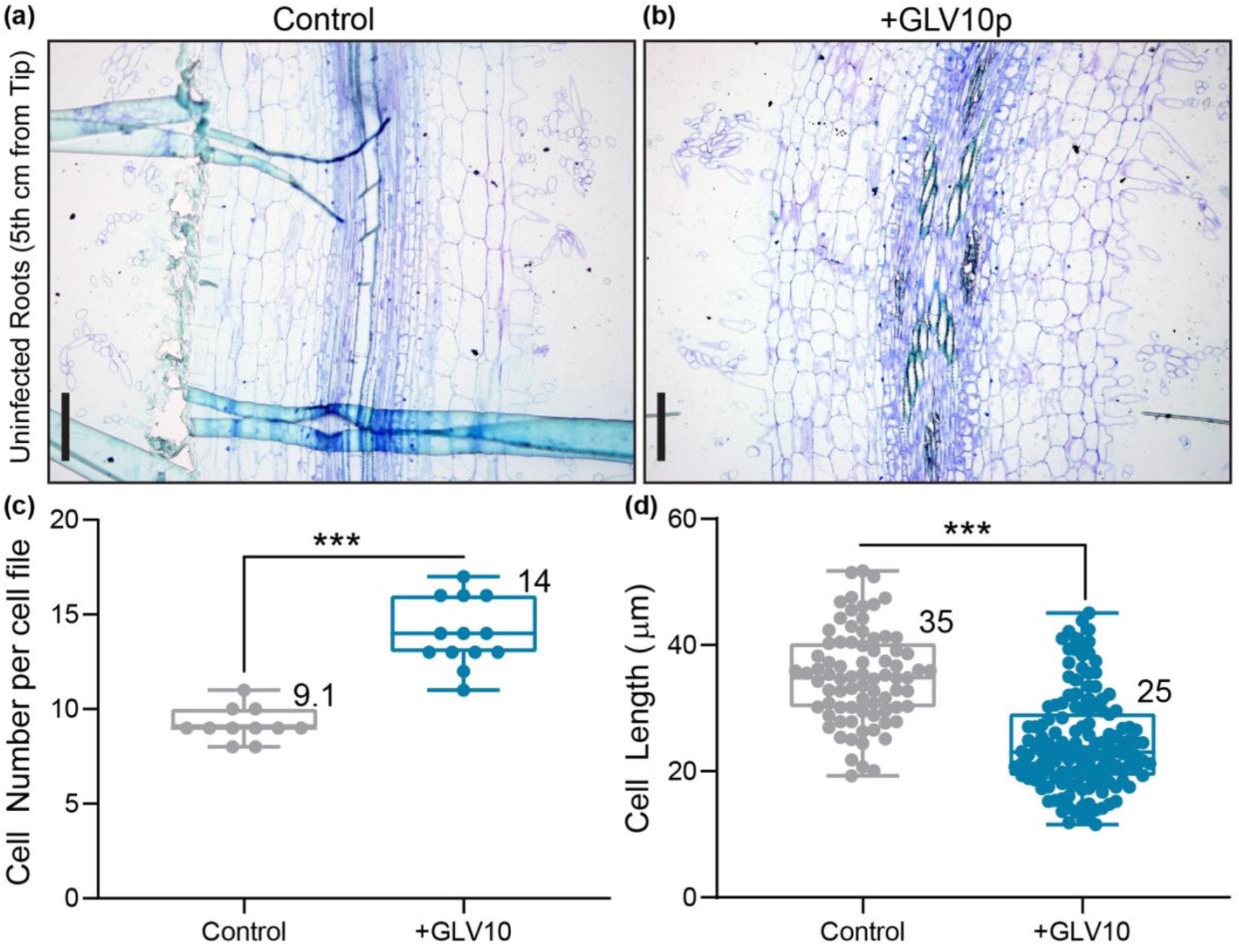
The synthetic peptide GLV10 affects cell number and cell size in Medicago roots. Images show 2.5 μm thick sections of root segments collected four cm below root tip at a 20x magnification. Control roots treated with solvent (a) and roots treated with 1 μM GLV10 peptide (b) for seven days. Scale bars represent 50 μm. Segments from at least eight roots per sample were analyzed. Quantification of data shown in (a,b) using ImageJ. Application of GLV10 peptide increases root length by increasing cell number (c) but decreasing cell size (d) in each cortical cell file. Roots were treated with peptide for seven days on plates and compared to untreated roots. Student’s t-test \*\*\**p*<0.001.

## Notes

### Competing Interest Statement

The authors have declared no competing interest.

